# The virome of New Zealand fur seals reveals novel viral diversity and evidence of continued Canine distemper virus circulation

**DOI:** 10.64898/2026.11.18.767762

**Authors:** Stephanie J Waller, Benjamin A White, Janelle R Wierenga, Alasdair A Hall, Moss Thompson, Andrew D Wilson, Harry S Taylor, Jody S Weir, Jemma L Geoghegan

**Affiliations:** Department of Microbiology and Immunology, Faculty of Biomedical Sciences, University of Otago, 720 Cumberland Street, Dunedin, 9016, New Zealand; Otago Peninsula Eco Restoration Alliance, 45 Pakihau Road, Dunedin 9077, New Zealand; Hall Marine Consulting, Wellington 6022, New Zealand; Department of Conservation, Te Papa Atawhai, 18–32 Manners Street, Wellington 6011, New Zealand; Animal Health Laboratory, Diagnostics, Readiness and Surveillance, Biosecurity New Zealand, Ministry for Primary Industries, Upper Hutt, 5018, New Zealand; Biosecurity, Surveillance & Incursion Investigation, Diagnostics, Readiness and Surveillance, Biosecurity New Zealand, Ministry for Primary Industries, 66 Ward Street, Upper Hutt, 5018, New Zealand

**Keywords:** Canine distemper virus, Marine mammal virome, New Zealand fur seal, Metatranscriptomics

## Abstract

Canine distemper virus (CDV) has recently been detected in New Zealand fur seals (*Arctocephalus forsteri*), including retrospective samples from 2020 and outbreaks associated with substantial mortality in late 2024 and 2025. However, the geographic distribution of CDV and the broader viral diversity harboured by New Zealand fur seals remain poorly understood. Characterising the fur seal virome can provide insights into viral diversity, dispersal and the occurrence of viral co-infections, while assessing the distribution of CDV across colonies. We used metatranscriptomic sequencing to characterise the viromes of 178 live and recently deceased fur seals sampled from four large fur seal colonies (Kaikōura, Dunedin, West Coast and Wairarapa). We investigated viral diversity, community composition and geographic structure using phylogenetic analyses to examine viral evolutionary relationships and CDV distribution. We identified 23 viral species, including 20 putatively novel species. Several were related to viruses previously identified in terrestrial mammals. One viral species was detected across geographically separated colonies, while several other closely related viruses were identified across colonies, with viral sharing most frequent between Kaikōura and Wairarapa. Viral diversity was similar between live and recently deceased fur seals, while community composition showed geographic structuring. CDV was detected only in Kaikōura, where the lineage associated with the 2024 outbreak remained in circulation. No CDV was detected in the other sampled colonies. This study reveals substantial previously undocumented viral diversity in New Zealand fur seals, improving our understanding of viral distribution and CDV circulation in marine mammal populations.

## Introduction

Members of the genus *Morbillivirus*, within the family *Paramyxoviridae*, are important pathogens of humans and other mammals, capable of causing severe disease and substantial mortality across a broad host range^1^. Morbilliviruses are enveloped viruses with non-segmented, negative-sense RNA genomes^1^. These viruses are known to infect immune and epithelial cells through the signalling lymphocyte activation molecule (SLAM/CD150)^2^ and nectin-4 receptors^3^, respectively, although other, as-yet-uncharacterised, receptors are also likely to play a role^1^. Infection can cause lymphoid depletion and immunosuppression, leaving affected individuals susceptible to secondary infections^1,4^. Morbilliviruses are highly contagious and are primarily transmitted through respiratory secretions and can result in large-scale epidemics and substantial mortality in susceptible populations^1,4^. Several members of the genus have had profound impacts on animal and human health, including measles virus, rinderpest virus and peste des petits ruminants virus^1,5^. Canine distemper virus (CDV), also known as canine morbillivirus, is particularly notable for its broad mammalian host range^1^. Although primarily associated with domestic dogs, CDV has been reported in more than 20 mammalian families across six taxonomic orders^6,7^, demonstrating its capacity to infect diverse hosts and cross species barriers.

CDV has also been associated with mass mortality events in phocids in the Northern Hemisphere, including killing more than 10,000 Caspian seals (*Phoca caspia*) in 2000 and thousands of Baikal seals (*Pusa sibirica*) in 1987^8,9^. These outbreaks highlight the potential consequences of morbillivirus infection, particularly in populations with limited prior exposure or reduced capacity to withstand substantial mortality. The broad host range of CDV further raises concerns about its potential to circulate among diverse wildlife species and establish infection in previously unaffected populations.

CDV has now been detected in New Zealand fur seals (*Arctocephalus forsteri*), including retrospective samples from a mortality event in Gore Bay in 2020 and large die-offs at Kaikōura in late 2024, and Kaikōura and Cape Palliser in 2025^10,11^. The origin of CDV in New Zealand fur seals remains unknown, as does the route by which the virus entered these populations. Its broad host range raises the possibility of cross-species transmission, potentially from dogs or ferrets^11,12^. Domestic dogs have historically been considered an important reservoir of CDV, while mustelids and other wildlife species are also susceptible to infection^1,12–14^. No naturally acquired CDV cases have been confirmed in New Zealand domestic dogs since 1983, although a small number of recent vaccine-associated cases have been reported involving strains distinct from that detected in New Zealand fur seals^15,16^. However, undetected infections in dogs, including those not presented for veterinary care, cannot be entirely excluded. The detection of CDV in New Zealand fur seals across multiple years and locations highlights the need to better understand its circulation and geographic distribution within this species. Despite the importance of these mortality events, the extent to which CDV occurs and is being maintained across New Zealand fur seal populations remains unclear.

Although New Zealand fur seals are currently classified as ‘Least Concern’, their populations remain below historical abundance following extensive eighteenth- and nineteenth-century sealing that brought the species close to extinction^17^. Despite legal protection, recovery varies among colonies, with ongoing threats including fisheries bycatch, environmental change and emerging disease^17–19^. Additionally, New Zealand fur seals are important marine predators that link food webs through their consumption of diverse prey, with their diet and foraging behaviour reflecting seasonal and regional variation in prey availability^20,21^.

The potential implications of widespread CDV infection may extend beyond fur seals to other native pinnipeds. The endemic New Zealand sea lions (*Phocarctos hookeri*) is classified as ‘Endangered’^22^ and although transmission of CDV from fur seals to sea lions has not been established in New Zealand, the broad host range of CDV and the overlapping ranges of these species makes cross-species transmission a relevant concern^11^. Understanding the distribution of CDV and the wider viral diversity circulating in fur seals is therefore important for informing disease surveillance and assessing potential risks to other vulnerable marine mammals.

Here, we used metatranscriptomic sequencing to characterise the viromes of 178 live and recently deceased New Zealand fur seals sampled from four regions across the North and South Islands of New Zealand: Kaikōura, Dunedin, West Coast and Wairarapa. We investigated viral diversity and community composition across locations and health status, examined the evolutionary relationships and geographic distribution of detected viruses, and assessed whether the CDV lineage associated with the 2024 Kaikōura outbreak was detected in other sampled colonies. This study expands our understanding of viral diversity in New Zealand fur seals.

## Methods

### Animal ethics and permits

All fieldwork and sampling were conducted in collaboration with the New Zealand Department of Conservation (DOC) as part of its routine wildlife management activities. As no additional procedures were undertaken specifically for this study, animal ethics and permits were not required.

### Sample collection

Samples were collected by DOC from New Zealand fur seal colony sites in the Dunedin, West Coast, Wairarapa, and Kaikōura regions (Figure 1). Samples were collected between August 2025 and January 2026. Two colony sites were sampled within the Dunedin region (Cape Saunders and Otago Peninsula Eco Restoration Alliance (OPERA) reserve Otago Peninsula), one each in West Coast (Cape Foulwind) and Wairarapa (Cape Palliser) regions, and three in the Kaikōura area (Kaikōura Peninsula, South Coast, and Ōhau Point). A total of 178 samples were taken from fur seals across these sites, with fresh faecal swabs taken from live fur seal colonies only when they could be confidently attributed to fur seals. Rectal swabs were collected from dead fur seals found in the Cape Palliser and Kaikōura areas, where recent CDV outbreaks had occurred. Although these deaths may have been associated with CDV, other causes, such as starvation, could not be excluded, as the cause of death could not be confirmed in this study. Samples were placed in tubes of DNA/RNA Shield (Zymo Research) and transferred to a −80°C freezer for storage until total RNA extraction.

**Figure 1.**
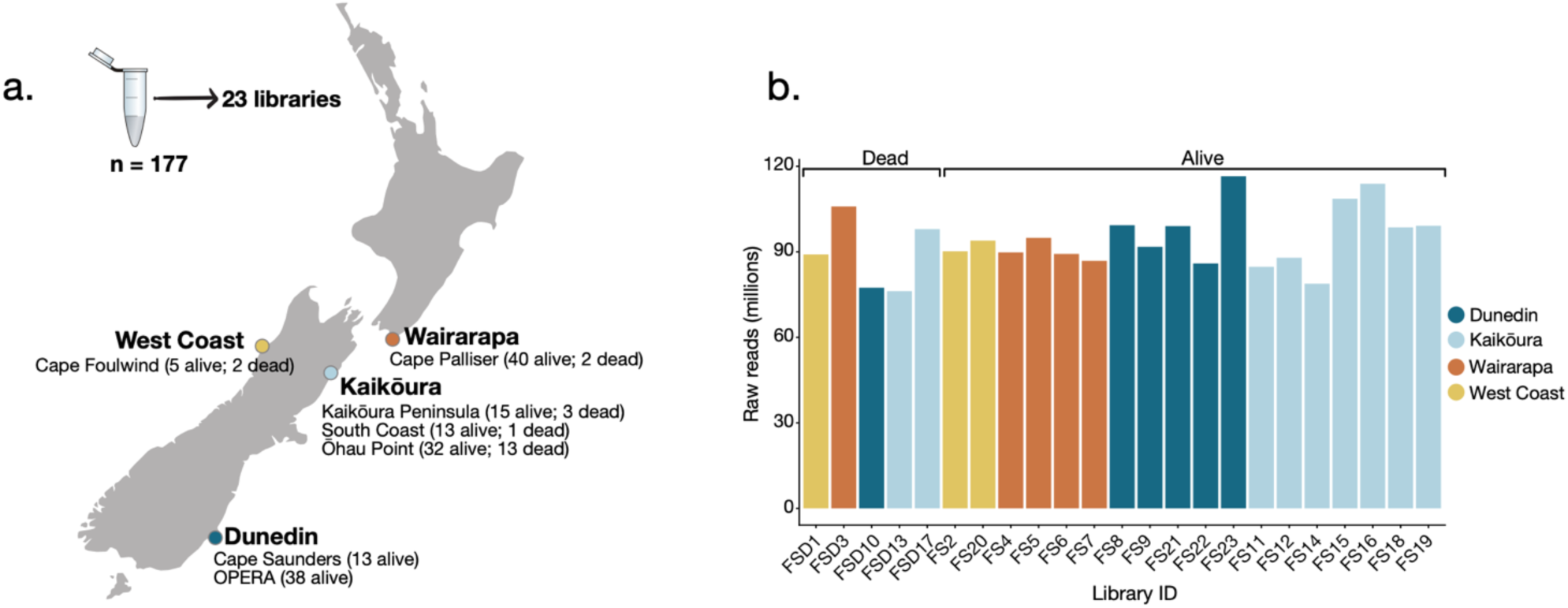
Sampling locations and sequencing read counts for New Zealand fur seals. (a) Map of New Zealand showing the seven sampling sites across four locations, with the number of samples collected from dead and live fur seals. In total, 177 samples were collected and pooled into 23 libraries. (b) Number of raw reads per library, with bars coloured by sampling region and libraries grouped by fur seal health status (dead or alive).

### Total RNA extraction and sequencing

Samples were thawed and removed from tubes using sterile tweezers, after which the swabs and 750μl of DNA/RNA Shield (Zymo Research) were transferred to ZR BashingBead Lysis Tubes. Tubes were shaken in a mini bead-beater 24 disrupter (Biospec Products) for five minutes to achieve cell lysis. Total RNA was extracted using the ZymoBIOMICS MagBead RNA kit (Zymo Research) following the manufacturers protocol. Extracted RNA concentration and quality were then measured via nanodrop spectrophotometer (Thermo Fisher), after which samples were stored at −80°C until pooling. Sample RNA was pooled based on location and fur seal health (Supplementary Table 1). One sample was excluded from RNA pooling and sequencing due to potential contamination with bird faeces observed during sample collection, leading to 177 samples across 23 RNA pools (Supplementary Table 1). Samples had an average RNA concentration of 30.89 ng/µL, while sample pools had an average RNA concentration of 29.5ng/µL (Supplementary Table 1).

Illumina Stranded Total RNA Prep (Illumina) was used in conjunction with Ribo-Zero Plus (Illumina) to generate complementary DNA libraries from pooled RNA, which were then amplified through 16 cycles of PCR. Total RNA sequencing was then conducted using Illumina NovaSeqX, generating 150bp paired-end reads with sufficient depth (>50 million reads per sample) to capture low abundance transcripts.

### Quality control, contig assembly and virus identification

Paired end adapter sequences (Nextera) and low-quality bases were trimmed using Trimmomatic (0.38)^23^. This was done using a four-base sliding window approach, where bases with a with a Phred score ≤ 5 were trimmed. Bases at the ends of reads were also trimmed until a quality score of three was reached, after which all reads shorter than 25bp were discarded, resulting in trimmed paired-end reads. Reads were then quality checked using FastQC (0.11.8)^24^ prior to de novo contig assembly using Megahit (1.2.9)^25^. Contigs were then searched against a local copy (downloaded 2024) of National Centre for Biotechnology Information’s (NCBI) protein database using Diamond BLASTx (2.1.9)^26^. Contig abundance was estimated using Bowtie2 (2.4.4)^27^. Contigs were sorted by kingdom using BLASTx’s ‘sskingdoms’ programme, where a maximum expected value of 1×10^−10^ for similarity was used as the cutoff threshold for putative viral contigs. Remaining non-viral hits were manually filtered, after which the remaining putative viral contigs were analysed using Geneious Prime (2026.0.2) to find, annotate, and translate potential open reading frames.

### Phylogenetic analysis of New Zealand fur seal viruses

Translated open reading frames of putative viral contigs were analysed using the NCBI standard protein BLAST online web search to identify viral RNA-dependent RNA polymerase or DNA polymerase encoding reads. These reads were aligned using the MAFFT (v. 7.490)^28,29^ L-INS-I algorithm, aligning reads against representative polymerase sequences from within the same viral family, as well as against their closest genetic matches in the NCBI GenBank database. Poorly aligned regions were removed with TrimAL (v.1.4.)^30^ using a gap threshold of 0.9. Phylogenetic trees were estimated using IQ-TREE (3.1.1)^31^, and annotated using Figtree (1.4.4)^32^.

### Bioinformatic analysis and interpretation of results

All statistical analysis and plots were generated using R (v4.6.1) and Rstudio (v2026.07.1 Build 147), in combination with the Tidyverse ggplot2 package^33^. Viral abundance for each family was standardised as reads per million, calculated by dividing the number of reads assigned to each family by the total number of reads in the corresponding library and multiplying by one million.

Alpha and beta diversity analyses were used to investigate the effects of sampling region and fur seal mortality status on viral diversity and community composition using the standardised viral abundance data. Alpha diversity was assessed using the Shannon index, which accounts for both viral family richness (the number of families detected) and evenness (the relative abundance of each family within a sample). Kruskal-Wallis testing was used to assess the significance of any differences in viral richness between groups. Additionally, for assessing differences in viral diversity between sampling regions, Pairwise Dunn’s testing combined with Benjamini-Hochberg correction was used to assess pairwise differences between all possible pairs of sampling regions whilst correcting for multiple comparisons.

Beta diversity was calculated using the Bray–Curtis dissimilarity index to quantify differences in viral community composition between samples using the vegan package (v2.7.5)^34^ in R (v4.6.1). Permutational multivariate analysis of variance (PERMANOVA) was used to test whether viral community composition differed according to sampling region and fur seal health status. Results were visualised using by generating ordination plots using non-metric multidimensional scaling based on the resulting Bray–Curtis dissimilarities, quantifying differences between viral communities on a scale from 0 (identical viral community compositions) to 1 (no shared viral families). Pairwise PERMANOVA testing, using Benjamini-Hochberg correction, was used to measure pairwise differences in viral community composition whilst correcting for multiple comparisons.

### Viral nomenclature

Viruses sharing <90% amino acid identity with their closest match to a conserved sequence (e.g., RNA-dependent RNA polymerase) within NCBI’s GenBank database were considered putatively novel^35^. Provisional names were assigned for each putatively novel species identified, each dependent on future validation by the International Committee on Taxonomy of Viruses. Viruses were named based on their host species (*Arctocephalus forsteri*) and the viral family they were closest to phylogenetically.

## Results

### Fur seal metatranscriptomes and raw read counts

We generated faecal and rectal swab metatranscriptomes from 177 New Zealand fur seals, comprising both live individuals and animals that had recently died, to investigate the prevalence of CDV and assess the occurrence of co-infections with other viruses (Figure 1). To this end, metatranscriptomic sequencing generated a total of 2.16 billion reads across all 23 libraries, with individual libraries yielding between 76 million and 117 million paired end reads (Figure 1, Supplementary Table 1). Libraries contained a mean of 93.7 million raw reads, providing substantial sequencing depth for downstream analysis of viral diversity and abundance.

### Differences in viral diversity across health statuses and sampling locations

We identified viral transcripts from a total of nine viral families across all libraries (Figure 2). Viruses from the *Sedoreoviridae* family made up a large majority (∼91-96%) of total virome diversity from both live and dead fur seal libraries (Figure 2a). Viruses from the *Picornaviridae* and *Paramyxoviridae* families appeared in greater abundance amongst dead fur seals, making up a combined 98.2% of all non-*Sedoreoviridae* viral reads in these samples. Comparatively, viruses from the *Caliciviridae* and *Astroviridae* families made up greater portions of total viral diversity in samples taken from live fur seals, making up 74.6% of non-*Sedoreoviridae* reads. Viruses from the *Nodaviridae* and *Phenuiviridae* families were only identified in libraries taken from live fur seals. Additionally, *Coronaviridae* reads were found in one library taken from dead fur seals. Notably, *Paramyxoviridae* made up a greater portion of viral diversity in libraries sampled from dead fur seals, accounting for 2.4% of total viral abundance in dead fur seals compared to 0.0131% of total abundance in live fur seals.

**Figure 2.**
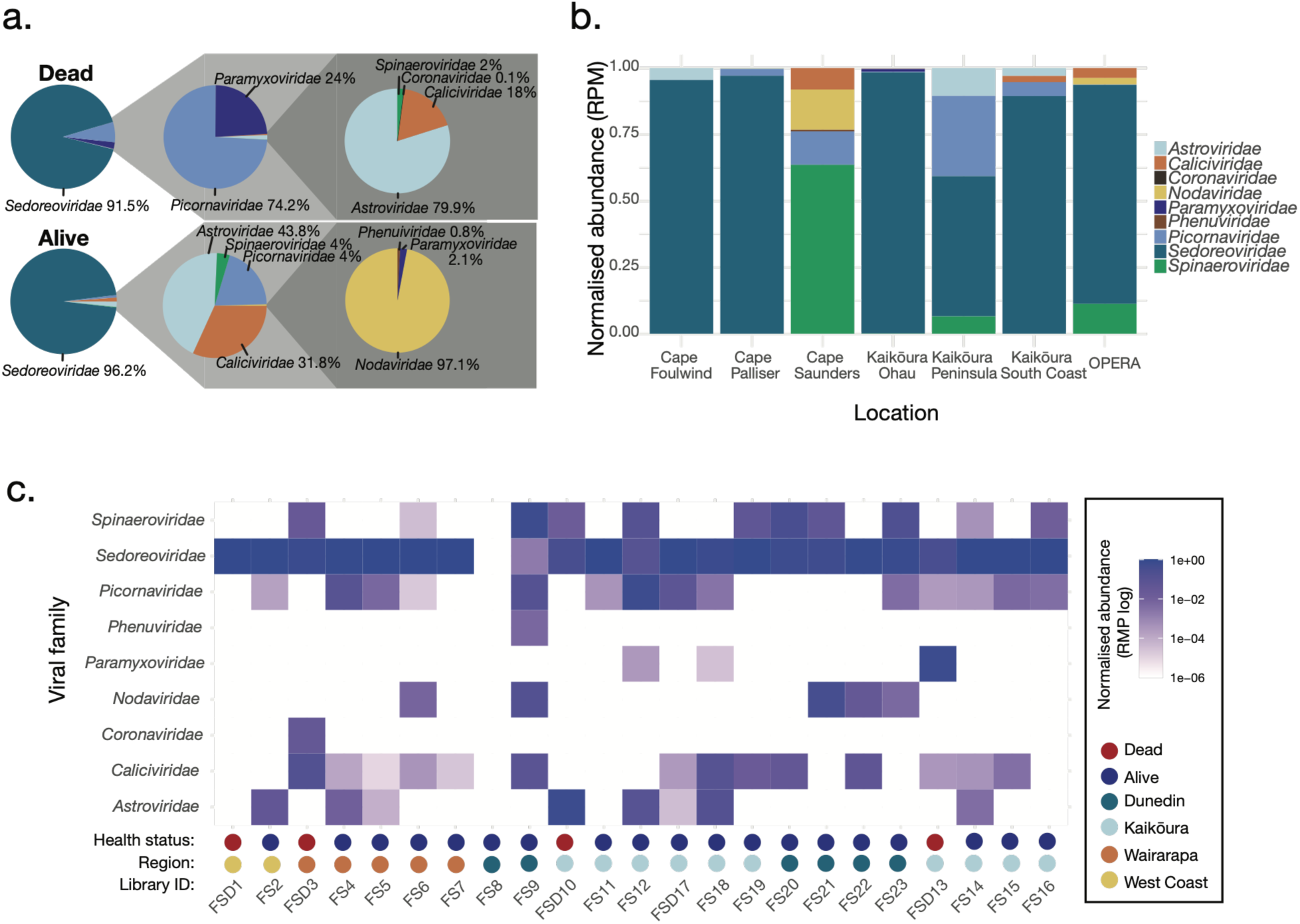
Viral community composition across fur seal libraries. (a) Pie charts showing the proportion of total viral reads assigned to each viral family in libraries from dead and live fur seals. (b) Stacked bar plot showing the normalised abundance of viral families across sampling locations, with colours indicating viral family. (c) Heat map showing normalised reads per million (RPM) for each viral family across fur seal libraries. The colour scale represents normalised abundance, with darker colours indicating higher abundance. Columns are ordered by sampling location and coloured by fur seal health status and sampling region.

When comparing viral diversity among sampling locations, *Sedoreoviridae* again made up the majority of viral reads across most sampling locations, although no *Sedoreoviridae* reads were detected from samples taken at Cape Saunders, Dunedin (Figure 2b). One library (FS8), taken from live fur seals at Cape Saunders, Dunedin, contained no reads corresponding to viral families known to infect mammals (Figure 2c). Of the nine viral families identified across the remaining 22 libraries, *Sedoreoviridae* were present in all of them, and were the most abundant viral family across most libraries. *Picornaviridae*, *Caliciviridae*, and *Spinaeroviridae* were the next most common, present in 15, 14, and 11 libraries, respectively, although not at consistently high abundances. Other families were considerably less abundant, with *Nodaviridae* only being found in five libraries and *Paramyxoviridae* only being found in three. Viruses from the *Coronaviridae* were only present in one library (FSD3) sampled from dead fur seals at Wairarapa, Cape Palliser and occurred at relatively low abundance (Figure 2c).

### Alpha and beta diversity analysis

To assess differences in viral diversity and community composition by sampling region and fur seal mortality status, we analysed alpha and beta diversity. Sampling region had no significant effect on alpha diversity where Shannon diversity did not differ significantly among regions (Kruskal–Wallis test, p = 0.26; Figure 3a). Since *Sedoreoviridae* accounted for a large proportion of viral reads across many samples, alpha diversity analysis was repeated after excluding this family to assess its influence on regional patterns. Results remained unchanged, with no significant differences in alpha diversity among regions (p = 0.32; Supplementary Figure 1a).

**Figure 3.**
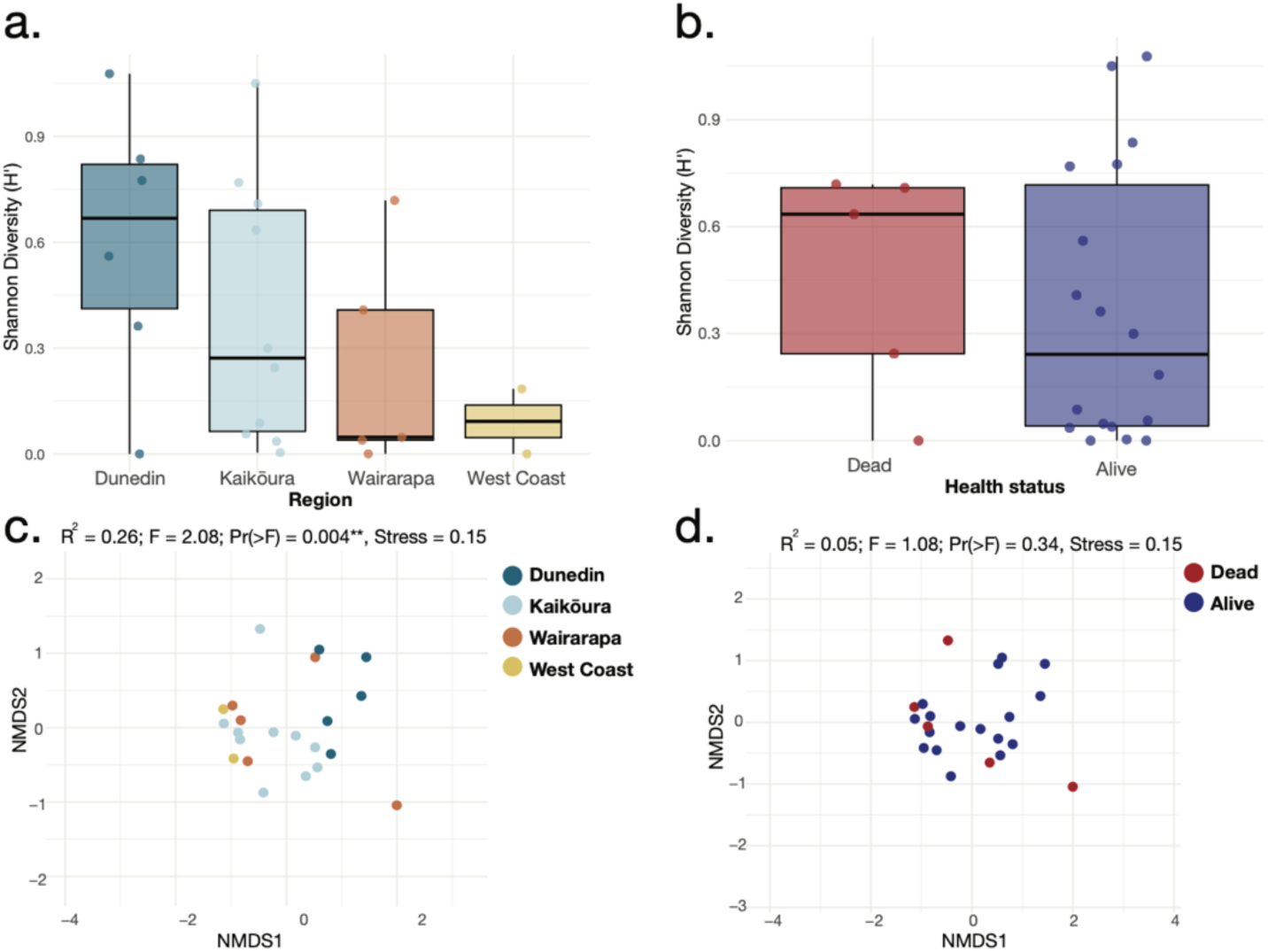
Full virome alpha and beta diversity analysis by region and dead/alive status. Shannon diversity plots, (a) by sampling region and (b) by whether fur seals were alive or dead. Non-metric multidimensional scaling plots depicting differences in virome composition, based on Bray–Curtis dissimilarities and tested using PERMANOVA, (c) by sampling region and (d) whether fur seals were alive or dead.

Beta diversity was used to assess differences in viral community composition among sampling regions. PERMANOVA showed that sampling region explained 25.7% of the variation in viral community composition (R² = 0.26, p = 0.004; Figure 3c). However, none of the pairwise regional comparisons remained statistically significant after Benjamini–Hochberg correction for multiple testing. Therefore, although sampling region was significantly associated with overall variation in viral community composition, no significant differences could be identified between individual pairs of regions.

Following the exclusion of *Sedoreoviridae*, sampling region remained significantly associated with viral community composition when all four regions were considered simultaneously, although the proportion of variation explained decreased from 25.7% to 19.4% (PERMANOVA, R² = 0.194, p = 0.031; Supplementary Figure 1c). As in the initial analysis, none of the pairwise comparisons between regions were statistically significant following Benjamini-Hochberg correction. These results indicate that the overall association between sampling region and viral community composition was not solely driven by the high abundance of *Sedoreoviridae*, although the reduction in the proportion of variation explained following their removal suggests that this viral family contributed to some of the observed regional variation.

Viral diversity and community composition were compared between live and deceased fur seals. No significant difference in alpha diversity was detected between groups based on Shannon diversity (p = 0.79; Figure 3b). Similarly, PERMANOVA found no significant association between fur seal status and viral community composition (R² = 0.05, p = 0.34; Figure 3d). To assess whether the high abundance of *Sedoreoviridae* influenced these results, analyses were repeated after excluding this family. Differences remained non-significant for both Shannon diversity (p = 0.22) and community composition (PERMANOVA, R² = 0.05, p = 0.31; Supplementary Figure 1d). Thus, no significant differences in viral diversity or community composition were detected between live and deceased fur seals, even after excluding *Sedoreoviridae*.

### Phylogenetic analysis of fur seal viruses

Phylogenetic trees were estimated to visualise the evolutionary relationships of identified viruses (Figure 4). Analyses were restricted to viral families for which assembled transcripts contained a polymerase gene (RNA-dependent RNA polymerase for RNA viruses or DNA polymerase for DNA viruses), as these conserved genes provide suitable markers for phylogenetic inference.

**Figure 4.**
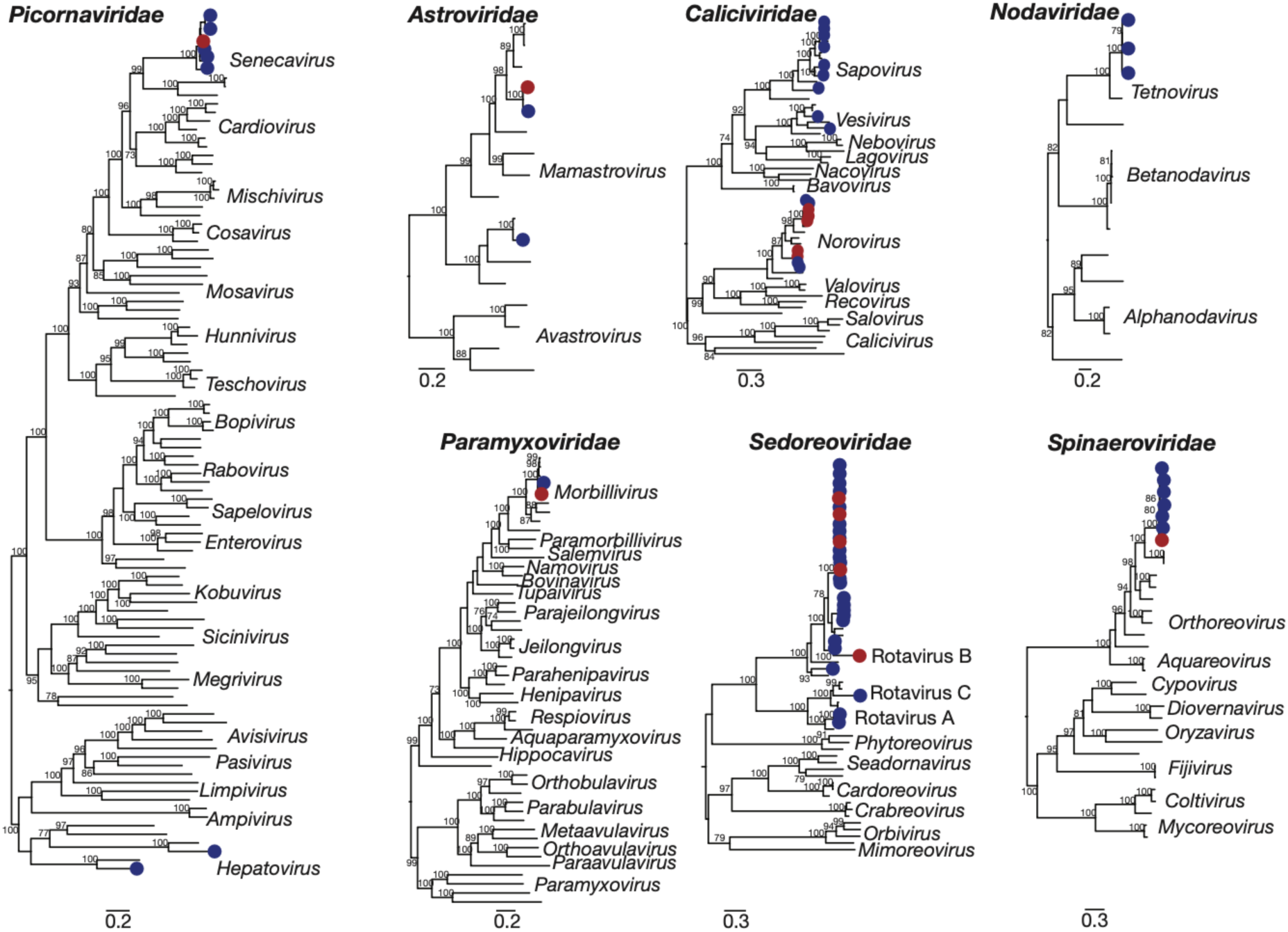
Maximum likelihood phylogenetic trees of RNA viruses found in fur seal viromes via metatranscriptomic sequencing. Trees were constructed using conserved RNA-dependent RNA polymerase sequence for seven viral families: *Picornaviridae*, *Astroviridae*, *Caliciviridae*, *Nodaviridae*, *Paramyxoviridae*, *Sedoreoviridae* and *Spinaeroviridae*. Trees were constructed to represent evolutionary relationships between identified viruses and closely related, previously characterised viruses, as well as their broader diversity. Identified viruses are displayed by dots, with blue representing sequences sampled from living fur seals, and red sampled from dead ones. Trees are rooted by midpoint with branch lengths corresponding to number of amino acid substitutions per site, indicated by scale bars. Branch nodes with bootstrap confidence intervals ≥70% are labelled.

*Coronaviridae* and *Phenuiviridae* were excluded because the assembled sequences did not contain sufficiently conserved gene regions for reliable phylogenetic analysis. All viruses identified clustered closely with established viral families and genera known to infect mammals. This phylogenetic analysis identified 23 viruses, of which 20 shared less than 90% amino acid similarity with their closest relative in NCBI’s GenBank database. These viruses were considered putatively novel and had names assigned. These names, as well as all information regarding the families, closest relatives, amino acid similarity percentages and sampling locations of the viruses identified, are provided in Supplementary Table 2.

In total we identified two viruses that shared greater than 90% amino acid identity with viruses previously detected in other marine mammals. These included Mamastrovirus 11 and California sea lion sapovirus 1, both previously identified in California sea lions (*Zalophus californianus*)^36,37^. Additionally, putatively novel sapoviruses, vesiviruses and noroviruses were identified that shared <90% amino acid identity to viruses that were previously identified in California sea lions^37–39^, while a putatively novel picornavirus was related to hepatovirus B1, previously reported in harbour seals^40^. All remaining viruses were most closely related to viruses characterised in terrestrial mammals, birds or reptiles, highlighting the diversity of viral lineages detected in New Zealand fur seals.

### Phylogenetic analysis of canine morbillivirus identified in fur seal viromes

We identified *Paramyxoviridae* sequences in our samples that clustered with members of the *Morbillivirus* genre (Figure 4 and Figure 5). Both were sampled from the Kaikōura region, with one sampled from live fur seals at South Coast (FS18) and the other sampled from dead fur seals at Ōhau point (FSD13). Neither sequence was considered novel, with the FS18 and FSD13 sequences sharing 96.09% and 95.47% amino acid identity with known CDV sequences, respectively. To assess the evolutionary relationship between these two viruses, a genus level phylogenetic tree was estimated aligning the identified viruses with representative morbilliviruses, as well as their closest genetic relatives available in GenBank (Figure 5). The two identified viruses clustered with other CDV viruses, and closest to the sequences sampled from the 2024 Kaikōura and 2025 Wairarapa, Cape Palliser CDV outbreak^10,11^.

**Figure 5.**
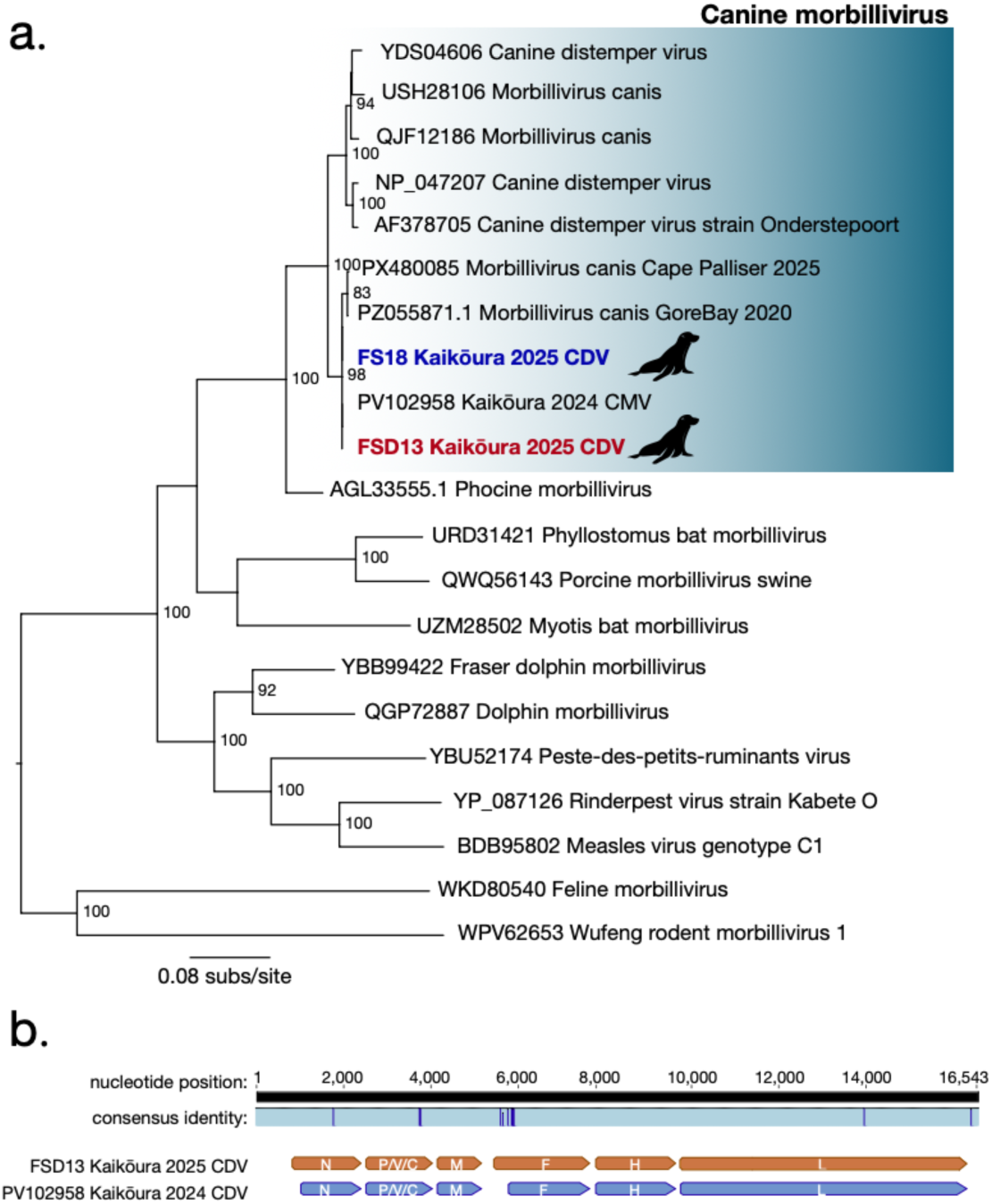
Phylogenetic analysis and sequence alignment of morbilliviruses found within Kaikōura fur seal viromes using metatranscriptomic sequencing. (a) Phylogenetic tree constructed using conserved RNA-dependent RNA polymerase sequence, to represent evolutionary relationships between the identified viruses and closely related, previously characterised viruses within the *Morbillivirus* genus. Identified viruses indicated in bold. Trees are rooted by midpoint with branch lengths corresponding to number of amino acid substitutions per site, indicated by scale bar. Branch nodes with bootstrap confidence intervals ≥70% are labelled. (b) Consensus identity genome alignment of identified CDV strain sampled at Ōhau point in 2025 and the 2024 Kaikōura CDV outbreak strain. Nucleotide differences are highlighted above open read frame annotations for the identified (orange) and reference (blue) strains.

The morbillivirus isolate identified in samples from dead fur seals at Ōhau Point (FSD13) comprised a complete, 16,543-nucleotide genome, compared with 15,392 nucleotides for the 2024 Kaikōura CDV outbreak strain^10^. Despite this difference in genome length, alignment of the two genomes revealed 99.84% nucleotide identity (Figure 5b).

## Discussion

This study provides a broad characterisation of the virome of New Zealand fur seals, identifying 23 viral species representing nine viral families. Viruses from seven families could be phylogenetically classified, while those from the remaining two families could not. Of the 23 viral species identified, 20 were considered putatively novel based on their divergence from previously characterised viruses. We also investigated the occurrence of viral co-infections, providing further insight into the diversity and potential interactions of viruses circulating within fur seal populations. These findings substantially expand current knowledge of viruses associated with New Zealand fur seals, which remain relatively understudied from a virological perspective^11,41,42^.

A key finding was the detection of CDV in two libraries from Kaikōura, including samples from live fur seals at South Coast and a recently deceased seal at Ōhau Point. The detection of a CDV lineage with 99.84% nucleotide identity to the 2024 Kaikōura outbreak strain suggests that this lineage has continued to be detected in New Zealand fur seal populations following the initial 2024 outbreak^10^. This is consistent with the ecology of CDV, which is often maintained and transmitted within densely populated animal groups where frequent close contact facilitates viral spread and persistence^43^. However, repeated introductions from an unsampled, non-fur seal, reservoir cannot be excluded.

The potential role of wildlife or domestic animal reservoirs, including mustelids and domestic dogs, warrants further investigation^12,44^. Similarly, limited information on fur seal movements and their interactions with other marine mammals, particularly beyond New Zealand waters^45,46^, means the potential for marine mammal cross-species CDV transmission remains largely unexplored. However, despite CDV detections in New Zealand fur seals, no confirmed mortality outbreaks due to CDV have been reported in marine mammals elsewhere in the Southern Hemisphere. CDV was not detected in samples from Wairarapa, West Coast or Dunedin, although this does not exclude infection in these populations, given potential influences of sampling effort, prevalence, timing and sample type. Future surveillance should prioritise areas of known CDV detection, particularly Kaikōura and Wairarapa, while incorporating potential terrestrial reservoirs, including dogs and mustelids. Broader genomic surveillance across wildlife populations will help determine whether this lineage is established in fur seal populations or is periodically introduced from an external source.

Although CDV has caused substantial mortality during outbreaks in pinnipeds^1^, detection of viral RNA in a pool of live fur seals indicates that infection is not necessarily associated with death at the time of detection. However, the subsequent clinical outcomes of these individuals are unknown. The detection of CDV in both live and recently deceased animals suggest that clinical outcomes may vary among individuals. Several factors could contribute to this variation, including co-infections with other pathogens, nutritional stress and starvation. CDV is known to induce immunosuppression through lymphocyte depletion and impaired immune function, increasing susceptibility to secondary infections^1,4^. Supporting the potential importance of secondary bacterial infections, a recent investigation of the 2025 CDV-associated mortality event in New Zealand fur seal pups isolated *Salmonella enterica* serovar Typhimurium from all seven deceased pups examined^11^. However, the contribution of this bacterium to disease severity and mortality remains unclear^11^. Although co-infections and nutritional status were not investigated in this study, their potential contribution to disease severity in New Zealand fur seals warrants further investigation. Future studies combining CDV screening with broader pathogen surveillance, nutritional assessments and clinical monitoring could help clarify the factors influencing infection outcomes in this population.

Beyond CDV, several identified viruses were related to those previously detected in other pinnipeds. These included Mamastrovirus 11 and California sea lion sapovirus 1, previously identified in California sea lions (*Zalophus californianus*), as well as a divergent vesivirus related to San Miguel sea lion virus 12 and a picornavirus related to hepatovirus B1, previously reported in harbour seals^36,37,40^. Such relationships may reflect shared viral lineages across marine mammal hosts, although the limited availability of pinniped viral sequences makes it difficult to distinguish broad geographic distributions from historical host associations or cross-species transmission. Expanded sampling of pinnipeds, particularly in the Southern Hemisphere, alongside comparative genomic analyses will help resolve these relationships.

Rotavirus-like viruses within the *Sedoreoviridae* were particularly prominent, with six putatively novel viruses identified and one detected across all four sampling regions. Rotaviruses have previously been characterised in several pinniped species, including New Zealand sea lions^37,47,48^, suggesting that they may be widespread components of pinniped viromes. However, their detection and abundance in faecal material do not establish active infection or pathogenicity. Individual-level testing, combined with clinical data and targeted investigation of animals with gastrointestinal disease, would help determine their significance for fur seal health.

Viral community composition was significantly associated with sampling region, although no individual pairwise comparisons remained significant after correction for multiple testing. This suggests that regional variation may occur across the dataset, but the available evidence does not establish clear differences between specific locations. Several viruses were detected across multiple regions, including a rotavirus found at all four locations. These patterns are consistent with geographically widespread viral lineages, potentially facilitated by fur seal movement or other transmission routes^45,46^. However, shared detection does not demonstrate direct transmission between colonies, and uneven sampling among regions may have influenced the observed patterns.

No significant differences in alpha diversity or overall viral community composition were detected between live and deceased fur seals. These findings remained unchanged after excluding the highly abundant *Sedoreoviridae*, suggesting that this family did not solely account for the observed patterns. However, mortality status is an imperfect proxy for disease outcome, as deceased fur seals may have died from causes unrelated to infection, or infection itself may have contributed indirectly to mortality through effects such as reduced feeding, weakness or impaired swimming ability^18,49,50^. Similarly, the clinical health of live fur seals was not established in detail. Consequently, the relationship between CDV infection and mortality cannot be established from detection status alone. Rather, overall virome composition did not clearly distinguish live from deceased animals within this study.

Several limitations should be considered when interpreting these findings. Sampling was uneven across locations and health-status groups, potentially reducing the power to detect regional differences or associations with mortality. In addition, faecal samples and rectal swabs primarily capture viruses present in or shed through the gastrointestinal tract and may not adequately represent viruses with respiratory or systemic tissue tropisms. Detection of viral sequences also does not confirm active infection, particularly for viruses that may originate from dietary or environmental sources, although the viruses in this analysis were thought to be mammalian-infecting. The cross-sectional sampling design provides a snapshot of viral diversity and cannot establish temporal patterns of infection or transmission. Finally, the limited number of available CDV genomes also prevents robust temporal analyses to distinguish between local persistence and repeated introductions, or to estimate when this lineage was introduced into New Zealand. Expanded genomic surveillance will be needed to resolve the evolutionary history and circulation patterns of this lineage.

Future studies should combine longitudinal and individual-level sampling with serology and clinical or necropsy data to distinguish active infection from incidental viral detection and investigate associations with disease. Continued CDV surveillance, including genome sequencing and investigation of potential terrestrial hosts, is a priority. Broader comparative sampling of New Zealand fur seals and other pinnipeds would also help clarify host associations and the geographic distribution of the newly identified viruses.

Overall, this study expands knowledge of the New Zealand fur seal virome and provides evidence of continued detection of a CDV lineage associated with disease in New Zealand fur seals, known to have been circulating since at least 2020. The identification of numerous divergent viruses highlights the extent of uncharacterised viral diversity in marine mammals and the value of continued genomic surveillance for understanding wildlife virology and monitoring pathogens of conservation concern.

## Resource availability

Viral sequences have been submitted to GenBank under the accession numbers QB356754-QB356825 while raw sequencing reads are available under BioProject PRJNA1554406.

## Acknowledgements

We would like to thank Kate Simister for her assistance with sample collection.

