## Supplementary Figure 1 for "The virome of New Zealand fur seals reveals novel viral diversity and evidence of continued Canine distemper virus circulation"

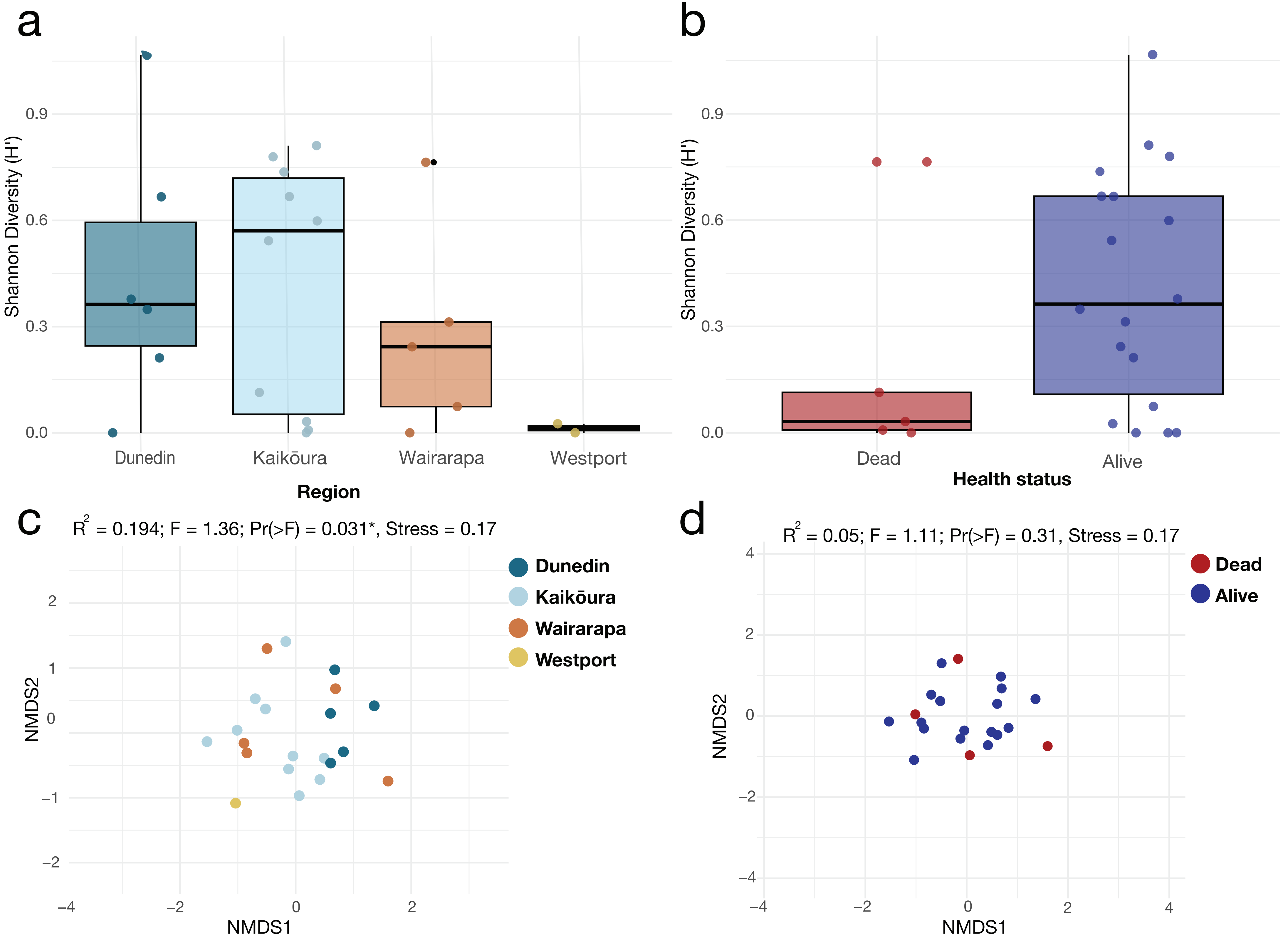

a
Shannon Diversity (H')
0.0
0.3
0.6
0.9
Dunedin
Kaikōura
Wairarapa
Westport
Region
b
Shannon Diversity (H')
0.0
0.3
0.6
0.9
Dead
Alive
Health status
c
R2 = 0.194; F = 1.36; Pr(>F) = 0.031*, Stress = 0.17
Dunedin
Kaikōura
Wairarapa
Westport
NMDS2
NMDS1
d
R2 = 0.05; F = 1.11; Pr(>F) = 0.31, Stress = 0.17
Dead
Alive
NMDS2
NMDS1
